# EasyEyes: Crowded Dynamic Fixation for Online Psychophysics

**DOI:** 10.1101/2025.02.26.640403

**Authors:** Fengping Hu, Joyce Yi Xin Chen, Denis G. Pelli, Jonathan Winawer

**Affiliations:** Department of Psychology, New York University, New York, NY, USA; Department of Biology, the University of Michigan – Ann Arbor, Ann Arbor, MI, USA; Center for Neural Science, New York University, New York, NY, USA

**Keywords:** fixation, online psychophysics, gaze tracking, EasyEyes, crowding, cursor tracking

## Abstract

Online vision testing enables efficient data collection from diverse participants but often requires accurate fixation. When needed, fixation accuracy is traditionally ensured by using a camera to track gaze. That works well in the lab, but tracking during online testing with a built-in webcam is not yet sufficiently precise. Kurzawski, Pombo, et al. (2023) introduced a fixation task that improves fixation through hand-eye coordination, requiring participants to track a moving crosshair with a mouse-controlled cursor. This *dynamic fixation* task greatly reduces peeking at peripheral targets relative to a stationary fixation task, but does not eliminate it. Here, we introduce a *crowded dynamic fixation* task that further enhances fixation by adding clutter around the fixation mark to leverage crowding. We assessed fixation accuracy during peripheral threshold measurement. Relative to the RMS gaze error during the stationary fixation task, dynamic fixation error was 61%, while crowded dynamic fixation error was only 47%. With a 1.5° tolerance, peeking occurred on 9% of trials with stationary fixation, 4% with dynamic fixation, and 0% with crowded dynamic fixation. This improvement eliminated implausibly low peripheral thresholds, likely by preventing peeking. We conclude that crowded dynamic fixation provides accurate gaze control for online testing.

## Introduction

Visual psychophysics often requires that the experimenter know where the participant is looking. Often, the participant is instructed to keep their eyes on a fixation mark. Ideally, the experimenter verifies fixation accuracy with a gaze tracker. This is especially important for eccentricity-dependent tasks, like acuity (Weymouth, 1958) and crowding (Bouma, 1970). Accurate gaze tracking, however, requires expensive equipment and in-lab testing. This excludes labs without the proper equipment and restricts the participant pool to the local population. This population is often WEIRD (Western, Educated, Industrialized, Rich, Democratic) and not representative of the global population (Henrich et al., 2010; Rad et al., 2018).

Online testing solves multiple problems. With no need for a lab, one can test a larger, more representative sample with far less effort. However, vision scientists have often missed out on this opportunity because of concerns about monitoring fixation. Several open-source packages are available for doing gaze tracking online using the computer’s webcam (Papoutsaki et al., 2016; Xu et al., 2015). However, software using the typical resolution of a built-in webcam limits the gaze-tracking accuracy to around 4 deg (Papoutsaki et al., 2016; Semmelmann & Weigelt, 2018). Recently, Kaduk et al. reported that adding a high-resolution webcam can decrease the gaze tracking error to 1.4 deg (Kaduk et al., 2024), but this requires buying the equipment for the participants, and frequent calibration. Alternatively, Kurzawski and colleagues (2023) showed that asking participants to use a cursor to track a moving fixation crosshair (“dynamic fixation”) yielded good fixation in most participants, meaning they kept their eyes fixed on the moving crosshair. We use the word “fixation” here to mean keeping the eyes on the crosshair, whether or not it is moving. The crosshair moves slowly and predictably, on a circular path, to make it easy to pursue. We wondered whether we can improve the method to achieve good fixation in all participants, eliminating the need for an eye tracker.

To understand our motivation, it helps to consider the fixation task together with the principal task of the experiment. Kurzawski et al. (2023) used the dynamic fixation task to measure a peripheral threshold. A trial begins with central fixation, followed by a brief flash of the stimulus, a three-letter triplet, in the periphery. Without an effort to suppress saccades, participants tend to move their eyes to the stimulus. We call this “peeking”. The stimulus is flashed briefly, to disappear before the participant’s eyes can reach it. Nonetheless, the participant knows a stimulus will appear, and might make an anticipatory eye movement, much as a soccer goalie sometimes dives to the left or right in anticipation of the penalty kick. Even if the eye movement only approaches the target by chance, it will sometimes land near the target, improving performance and corrupting the threshold estimate (Kurzawski, Burchell, et al., 2023).

The participant feels the tension of opposing attractions to gaze. On the one hand, the dynamic fixation task encourages fixating the crosshair because, when tracking an object with your hand, it is natural to track with your eyes as well (Koken & Erkelens, 1992; Xia & Barnes, 1999). On the other hand, it is also natural to shift gaze toward an anticipated target. The current study sought to enhance the tendency to fixate, so it would always win.

To assess progress toward this goal, we compared three fixation tasks: stationary fixation, dynamic fixation, and crowded dynamic fixation. In stationary fixation, a static crosshair is displayed at the center of the screen, serving as a visual anchor for the participant’s gaze. This is the traditional way to present the fixation mark in psychophysical testing with peripheral stimuli. In dynamic fixation (Kurzawski, Pombo, et al., 2023), participants are asked to use the mouse and mouse-controlled cursor to manually track a crosshair moving along a circular trajectory. In *crowded dynamic fixation*, the tracking task is enhanced by the addition of three distractors moving randomly near the moving crosshair. We implemented all three kinds of fixation while doing a peripheral threshold task and used a gaze tracker to measure fixation accuracy.

Our results validate the effectiveness of crowded dynamic fixation in maintaining stable central fixation during peripheral visual testing. Among the three types of fixation task, crowded dynamic fixation is the most effective in keeping participants’ gaze at fixation. Crowded dynamic fixation provides this benefit with negligible increase in the experiment duration. The new crowded dynamic fixation task enables online testing with accurate fixation.

## Methods

### Participants

Twenty-one participants, aged 17 to 41, did our experiment. One participant was excluded because the Eyelink 1000 could not reliably track their eyes, and two participants were excluded from the crowding threshold analysis because of a software failure on that day. All participants were native English speakers, and, if they needed optical correction, they wore contacts. None of the participants had prior experience with psychophysical testing. Participants aged eighteen and older signed consent and debriefing forms. Participants under eighteen years old signed assent and debriefing forms, and their parents or guardians signed consent forms. All participants were compensated $15/hour for their participation. This experiment was approved by the New York University Institutional Review Board (IRB-FY2016-404).

### Apparatus

The EyeLink 1000 eye tracker (SR Research, Ottawa, Ontario, Canada) with a tower-mount setup was used to track participants’ right eye throughout the experiment at a sampling frequency of 1 kHz. An Apple iMac 27″ computer presented stimuli on an external monitor, LG 27” UltraFine 5K (27MD5KL-B), with a 59.5 × 33.5 cm screen with 5,120 × 2,880 pixels, and a frame rate of 60 Hz, and a luminance of about 275 cd/m^2^. The participant placed their chin on an adjustable chin rest 40 cm away from the monitor, and used a mouse to perform the tasks.

### EasyEyes

We used EasyEyes, an open-access web app for creating and running psychophysical experiments (Jiang et al., 2021). EasyEyes experiments are created by uploading a Microsoft Excel file, with one column per condition, specifying the desired parameters (such as stimulus size and duration). EasyEyes calibrates the screen size by asking the participant to indicate the size of a familiar object (such as a credit card) on the screen (Li et al., 2020). EasyEyes uses the PsychoJS implementation of Quest (*PsychoJS*, n.d.; Watson & Pelli, 1983) to measure thresholds. It records the location of the participant’s cursor and the fixation crosshair throughout the experiment at a frequency of 60 Hz. Gaze tracking data is recorded separately and integrated at the end of the experiment (see Data Analysis).

### Fixation task

Participants were asked to keep the cursor on the crosshair, which was either static or moving (described below). The cursor is “on” the crosshair whenever the cursor tip is inside a 0.15 deg “hotspot radius” of the crosshair center. Cursor tracking feedback is provided by increasing the crosshair stroke thickness 1.5-fold to indicate that the cursor is on the crosshair. The 0.15-deg hotspot radius was chosen based on pilot experiments that included cursor tracking and gaze tracking to minimize cursor and gaze errors, and to limit the perceived difficulty (Supplementary Figure 1).

#### Stationary fixation

A static crosshair is displayed at the center of the screen (Figure 1a). The cross consists of a horizontal and a vertical black line, each 1 degree in length and 0.05 degrees thick.

**Figure 1.**
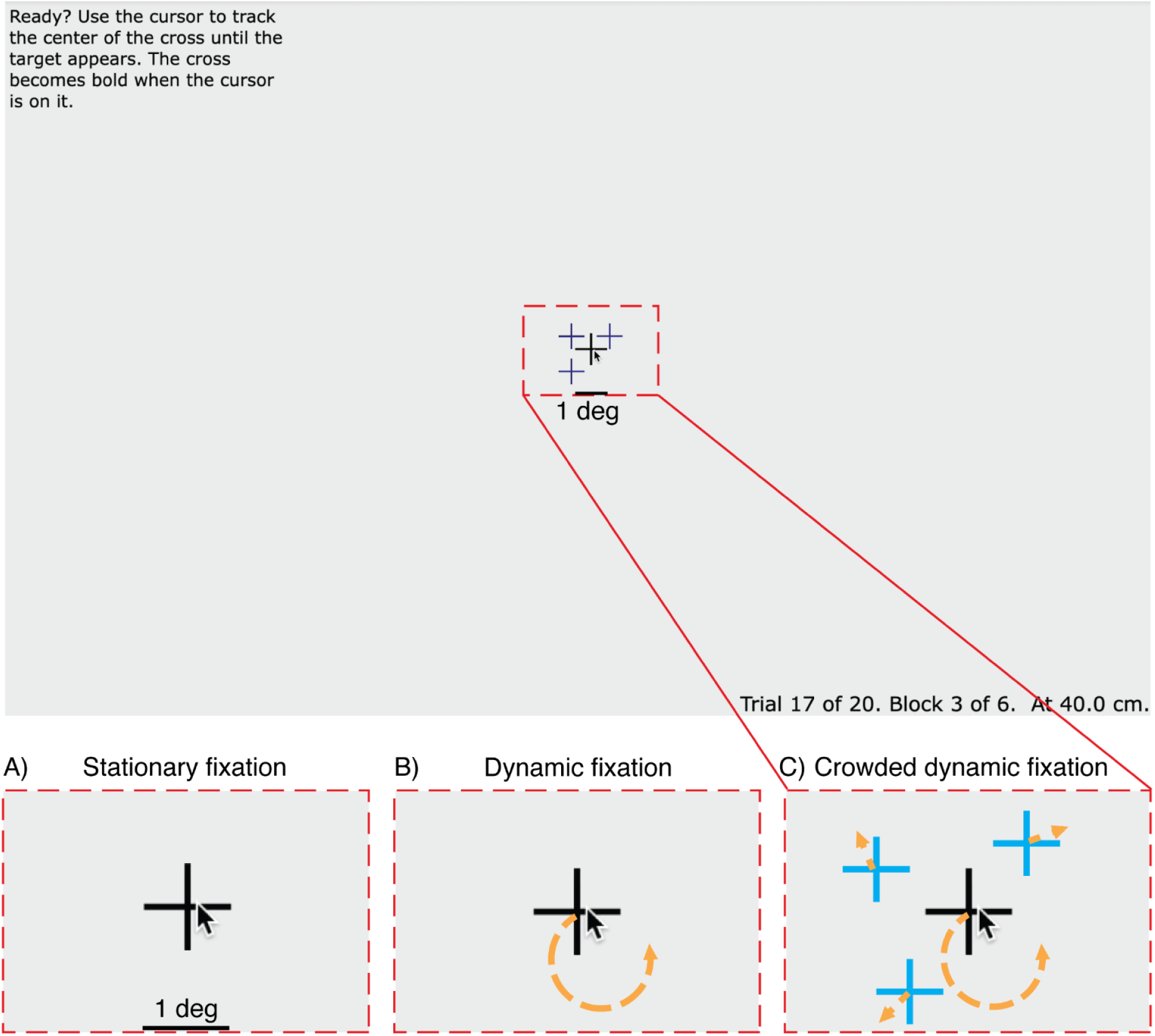
Tracking Task. Participants are asked to keep the cursor on the crosshair. The red dashed rectangle indicates the size of the area depicted in A-C. A) *Stationary fixation*: The crosshair is static. B) *Dynamic fixation*: The crosshair moves along a circular trajectory at a constant speed (indicated by the orange arrow). C) *Crowded dynamic fixation*: The moving crosshair (black) is surrounded by three distractors. For display purposes, the color difference between fixation and distractors is exaggerated.

#### Dynamic fixation

The crosshair (same as above) moves counterclockwise at a speed of 0.4 deg/sec along a circular trajectory. The trajectory centers at the center of the screen and has a radius of 0.5 deg (Figure 1b). Each trial begins with the crosshair at a random point on this trajectory. The motion speed and motion radius of the crosshair were chosen to minimize tracking error, gaze error, and perceived difficulty (Supplementary Figure 1).

#### Crowded dynamic fixation

The crosshair motion is the same as for dynamic fixation, but three distractors are added near the crosshair (Figure 1c). The distractors were added to make it difficult to see the crosshair when looking away (i.e., breaking fixation) but not when fixating it. Each distractor consists of a horizontal and a vertical dark blue line with a length of 0.8 deg and a thickness of 0.05 deg. Thus the distractor crosshairs bear a family resemblance to the fixation crosshair, but have obviously different color and size. The three distractors move in Brownian motion with an instantaneous speed of 1.1 deg/sec within 1 deg of the crosshair. To avoid clustering, a repulsive displacement is added to each distractor. The repulsion is implemented using the standard gravitational formula but with repulsion instead of attraction. We compute a repulsion direction and length from each pair and sum these to determine the displacement of each distractor. The direction of displacement for each pair is in the direction of the line connecting the pair, away from the other distractor. The length *h* of the displacement (in deg) between distractors *i* and *j* is:

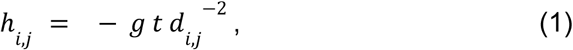

where *t* is the frame duration (sec), *d_i,j_* is the distance (deg) between distractors *i* and *j*, and *g* is 0.25 deg^3^/s.

### Crowding task

Each trial consists of three intervals: Track, View, and Respond (Figure 2). During the “Track” interval, participants are instructed to keep the cursor tip on the crosshair center. The tracking task continues until the participant has tracked continuously for the required duration (0.75 to 1.25 sec, randomly selected for each trial). If tracking breaks (cursor leaves hotspot radius), then the trial resets and waits for tracking to resume, when a new random-length interval is sampled. This tracking and resetting continues until the participant manages to track for a full (random length) interval. If a participant fails to initiate a trial after 30 seconds of tracking, the trial is skipped.

**Figure 2.**
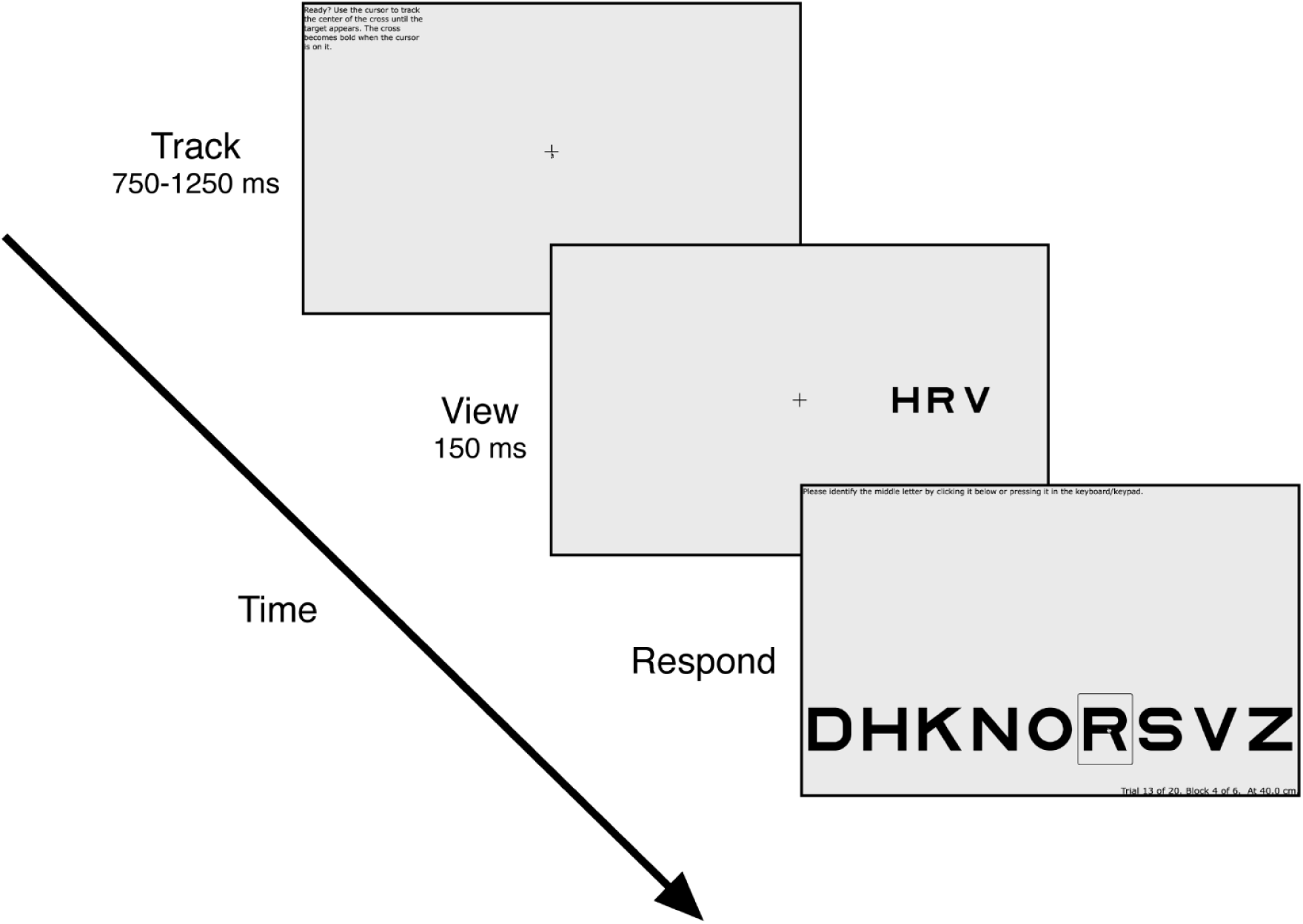
Crowding Experiment Procedure. *(1) Track:* During each trial, the participant is first asked to use a mouse to make the mouse-controlled cursor track the fixation crosshair. *(2) View:* After continuous successful tracking for 750 to 1250 ms (random from trial to trial), a letter trigram is shown for 150 ms, centered ±10 deg from fixation (i.e. current crosshair location), on the horizontal meridian. The crosshair remains on the screen at its final position. *(3) Respond:* After 150 ms, the trigram disappears, and the participant identifies the target letter by pressing it on the keyboard or clicking one of the nine letters along the bottom of the screen.

Upon completion of tracking, the trial proceeds to the “View” interval. The crosshair remains frozen on the screen at its final position, while the cursor and any distractors disappear. Simultaneously, a letter trigram appears for 150 ms, centered at ±10 deg eccentricity (relative to the current position of the fixation crosshair). The position of the trigram (left or right meridian) is random across trials. The size of the letters is 0.71 times the center-to-center spacing. Size and spacing co-vary across trials, controlled by the staircase. The three letters in the trigram are chosen randomly, without replacement, from nine letters, DHKNORSVZ, of the Sloan font (Pelli et al., 1988; Sloan, 1959). The letter C was excluded because it is too easily confused with O (Elliott et al., 1990). The middle letter is the target.

The “Respond” interval begins after the 150-ms “View”. The crosshair disappears and an array of all 9 possible target letters appears at the bottom of the screen. The participant is asked to identify the target by pressing that key in the keyboard or clicking that letter on the screen.

### Experimental procedure

Each participant completed two in-lab sessions on separate days. The first session consisted of a practice experiment and the first run of the full experiment. The second session was the second run of the full experiment.

In the practice experiment, participants completed 6 blocks of 20 trials each. Each trial followed the general procedure of the crowding task described above, including the random left vs. right position of the target on each trial. However, for the first three practice blocks, only a single letter, rather than the letter trigram, was presented during the “View” interval. Participants were instructed to select the letter they saw during the “Respond” interval. Fixation was stationary in the first block, dynamic in the second, and crowded dynamic in the third. These initial blocks were designed to familiarize participants with the general experimental procedure and the three fixation tasks. The final three practice blocks were the same except that they included the letter trigrams.

The main experiment began with calibrating the eye tracker. The experimental session consisted of 3 blocks of 70 trials each. Within each block, the fixation condition was constant, either stationary, dynamic, or crowded dynamic, with the order randomized. Each 70-trial block consisted of two randomly interleaved 35-trial staircases, one for right and one for left. The center-to-center letter spacing was controlled by QUEST (Watson & Pelli, 1983). The second session was identical to the first, except for no practice and with new random ordering of blocks. At the end of each block, QUEST estimates a threshold for each condition (left or right) as the 53rd quantile of the posterior probability density function of threshold versus spacing.

### Data analysis

#### EasyEyes-Eyelink time alignment

EasyEyes records crosshair and cursor position (in each frame, at 60 Hz), while a MATLAB program records the gaze position (at 100 Hz) measured by EyeLink. Both programs save the data as CSV files including POSIX timestamps. The POSIX timestamps are used to align the two data files.

#### Gaze correction

Despite calibration of the eye tracker before the experiment, there is some drift in calibration over the course of the experiment. For each trial, we correct for this error by calculating the average gaze position in the last 400 ms of the “Track” interval and subtracting that from the measured gaze position in every frame of that trial.

To remove blinks from the eye tracking data, we first identify blinks by calculating the speed and velocity of gaze. A blink is detected when the velocity exceeds 1000 deg/sec or the gaze position deviates more than 20 degrees from fixation (likely off-screen). The start and end frames of each blink are then marked, and the entire blink period, along with an additional 100 ms before and after, is replaced with the eye position recorded 100 ms after the blink. If a blink occurs at the end of a trial, the blink period is replaced with the eye position recorded in the frame 100 ms before the blink.

## Results

Crowding thresholds were measured at ±10 deg eccentricity on the horizontal meridian using each of three fixation tasks: stationary, dynamic, and crowded dynamic. The crowding triplet appeared randomly on either the left or the right meridian. This was implemented by randomly interleaving two staircases, one for left and another for right. Gaze position was monitored by an eye tracker.

### Crowded dynamic fixation reduces gaze deviation throughout the trial

We first demonstrate the advantage of the crowded dynamic fixation task by examining gaze deviation at each time point in a trial. We do so by plotting gaze position during every trial, grouped by fixation condition, for an example participant (Figure 3). The traces indicate the horizontal (left column) or vertical (right column) distance between crosshair and gaze. The meridian of the crowding triplet is indicated by color (red for right, green for left).

**Figure 3.**
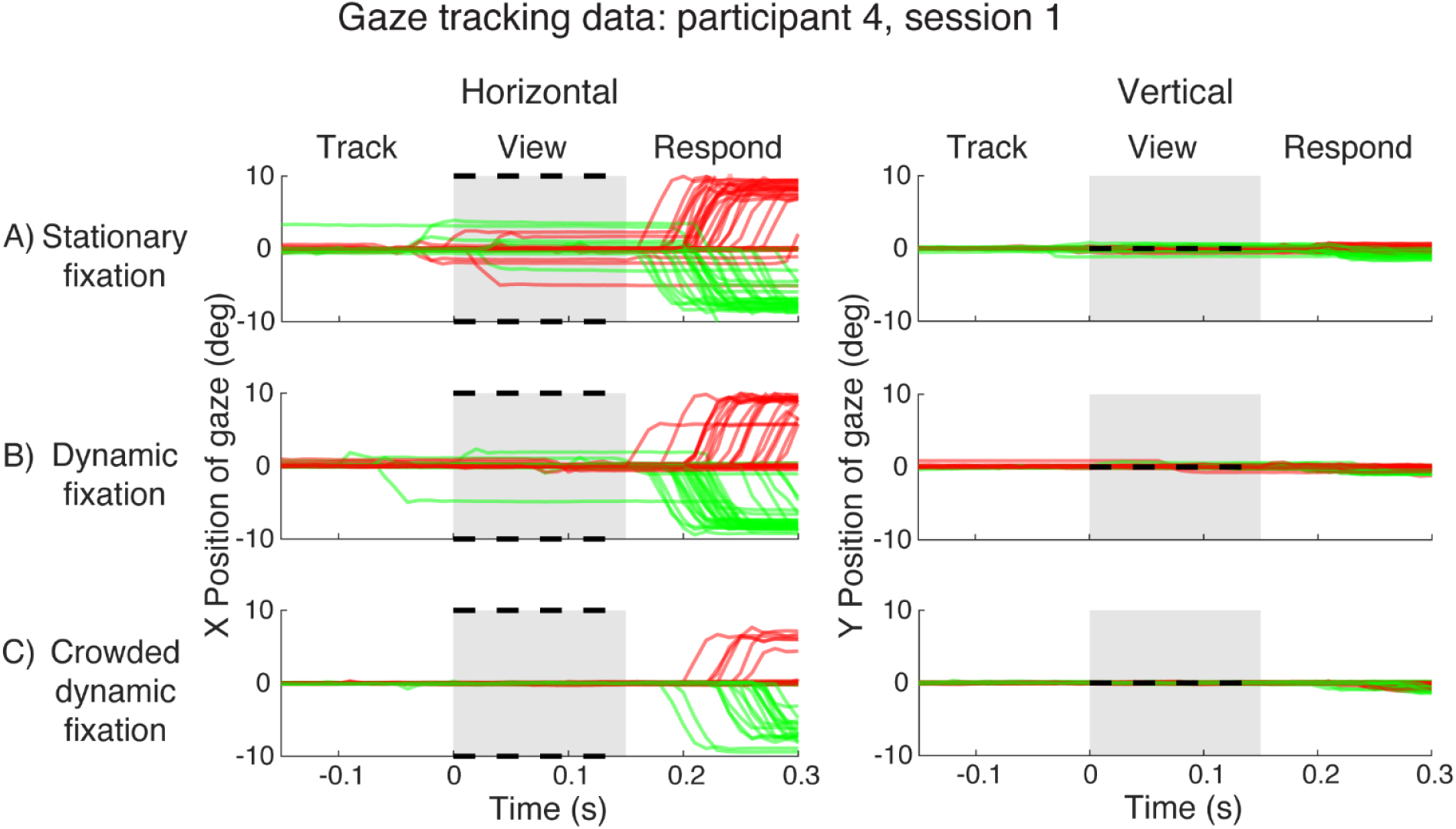
Gaze data for participant 4. A) Horizontal and vertical gaze position, relative to the crosshair position, during stationary fixation trials. The gray patch indicates target presence on the screen. Black dotted lines mark the two possible target locations (±10° horizontally). Each red or green trace is gaze position over time during one trial. Red traces are for trials with right targets and green traces are for trials with left targets. B) Gaze position during dynamic fixation trials. Plotted gaze position is relative to current crosshair position. C) Gaze position during crowded dynamic fixation trials. *(plot_all_trials_xy.m)*

With stationary fixation (Figure 3A), this participant made saccades (i.e. eye movements ≥ 1 deg) during 10 of 70 trials in either the “Track” or “View” interval. These saccades are problematic for measuring peripheral thresholds because they change the target eccentricity from the nominal 10°. These saccades were always on the horizontal axis, typically only a few degrees, much less than the 10° stimulus eccentricity, and in a direction (left vs right) uncorrelated with the unpredictable left-or-right target location. The timing, amplitude, and direction of the saccades indicate that the participant had some expectation of *when*, though not *on which side*, the stimulus would appear. We refer to saccades initiated during the Track or View interval as “*anticipatory saccades*”. In contrast, during the “Respond” interval, the participant usually made a saccade to the target location (10° amplitude, correct side). These saccades were clearly in response to seeing the target, and inconsequential because the stimulus disappeared before the saccade began. We refer to these as “*response saccades*”. A key goal of this project is to minimize anticipatory saccades. Response saccades don’t matter The frequency and amplitude of anticipatory saccades decreased with dynamic fixation, as reported by (Kurzawski, Pombo, et al., 2023). However, there were still four anticipatory saccades. The crowded dynamic fixation eliminated anticipatory saccades. There were no vertical saccades in any of the three fixation conditions.

Improvement in fixation accuracy from stationary to dynamic to crowded dynamic is shown by summarizing fixation accuracy at each time point in the trial. Specifically, we calculated the root mean square distance between the crosshair (i.e., fixation mark) and gaze (RMSE), across the 70 trials per fixation condition, separately for each time point (video frame) within the trial. This summary metric is shown for one participant (Figure 4A) and the group average (Figure 4B). The important part of the trial is the View interval, when the stimulus is on the screen (shaded gray region). During the View interval, the RMSE is highest for stationary fixation, intermediate for dynamic fixation, and lowest for crowded dynamic fixation, for both the example participant (Figure 4A) and the group average (Figure 4B). The data are replotted with RMSE calculated relative to stationary fixation in panels 4C and 4D. For the group average, the RMSE for dynamic fixation is about 60% and crowded dynamic fixation is about 45%, relative to stationary fixation.

**Figure 4.**
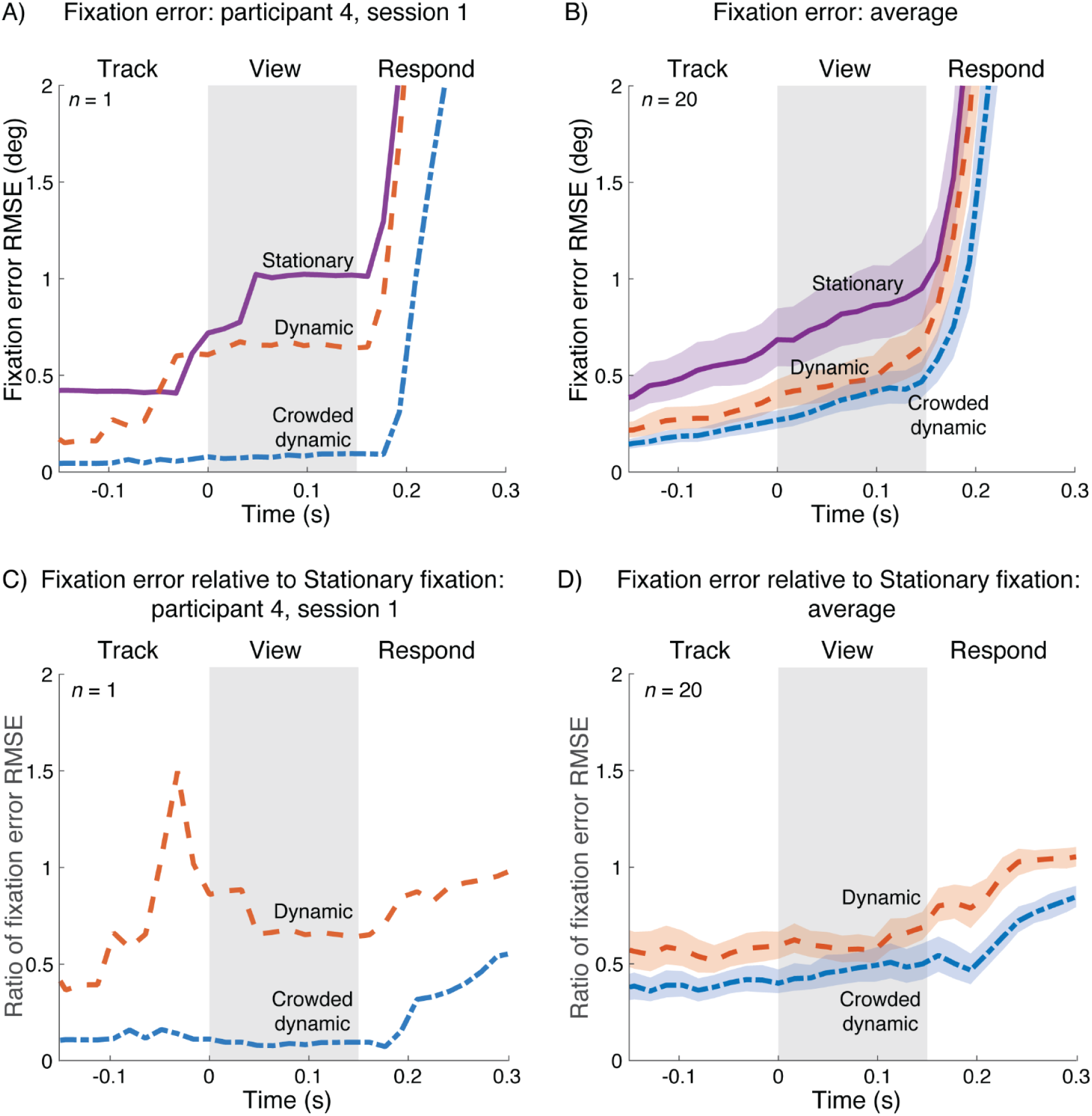
Fixation error for three fixation tasks. A) Distance between gaze position and crosshair position for participant 4, session 1. Each curve represents the RMSE over the 70 trials for that condition at each time point. *(plot_rmse_single_participant.m)*; B) Geometric means of fixation error RMSE from 40 experiment sessions (20 participants, 2 sessions each). *(plot_rmse_over_time_group.m, bootstrap_plot_rmse_group.m)*; Panels C,D: Same as A and B, but relative to the RMSE fixation error during stationary fixation (*plot_rmse_single_participant_ratio.m*, *bootstrap_plot_rmse_group_ratio.m*). For panels B and D, shaded regions indicate 68% confidence intervals bootstrapped across participants.

### Crowded dynamic fixation reduces the number of trials with large eye movements

The previous section reported the benefit of crowded dynamic fixation using a continuous measure (RMSE gaze deviation over time). Here we quantify the benefit with a discrete measure, the number of trials with large gaze deviations, a common metric of gaze behavior for psychophysical experiments.

Crowded dynamic fixation greatly reduced the number of trials with large gaze deviations, or “*fixation breaks*”. We calculated the number of such trials using several different tolerances for a fixation break (Figure 5). The trial is classified as having broken fixation if the participant’s fixation error (gaze distance from crosshair) exceeds tolerance at any time during the “View” interval. With 1.5 deg tolerance, fixation breaks are reduced from 9% (6 of 70 trials) with stationary fixation to 3% (2 of 70) with dynamic fixation, to none with crowded dynamic fixation. This reduction from stationary to dynamic is comparable to the results reported by Kurzawski et al. (2023). The order is the same at every tolerance: largest number of fixation breaks for stationary fixation, fewest for crowded dynamic fixation.

**Figure 5.**
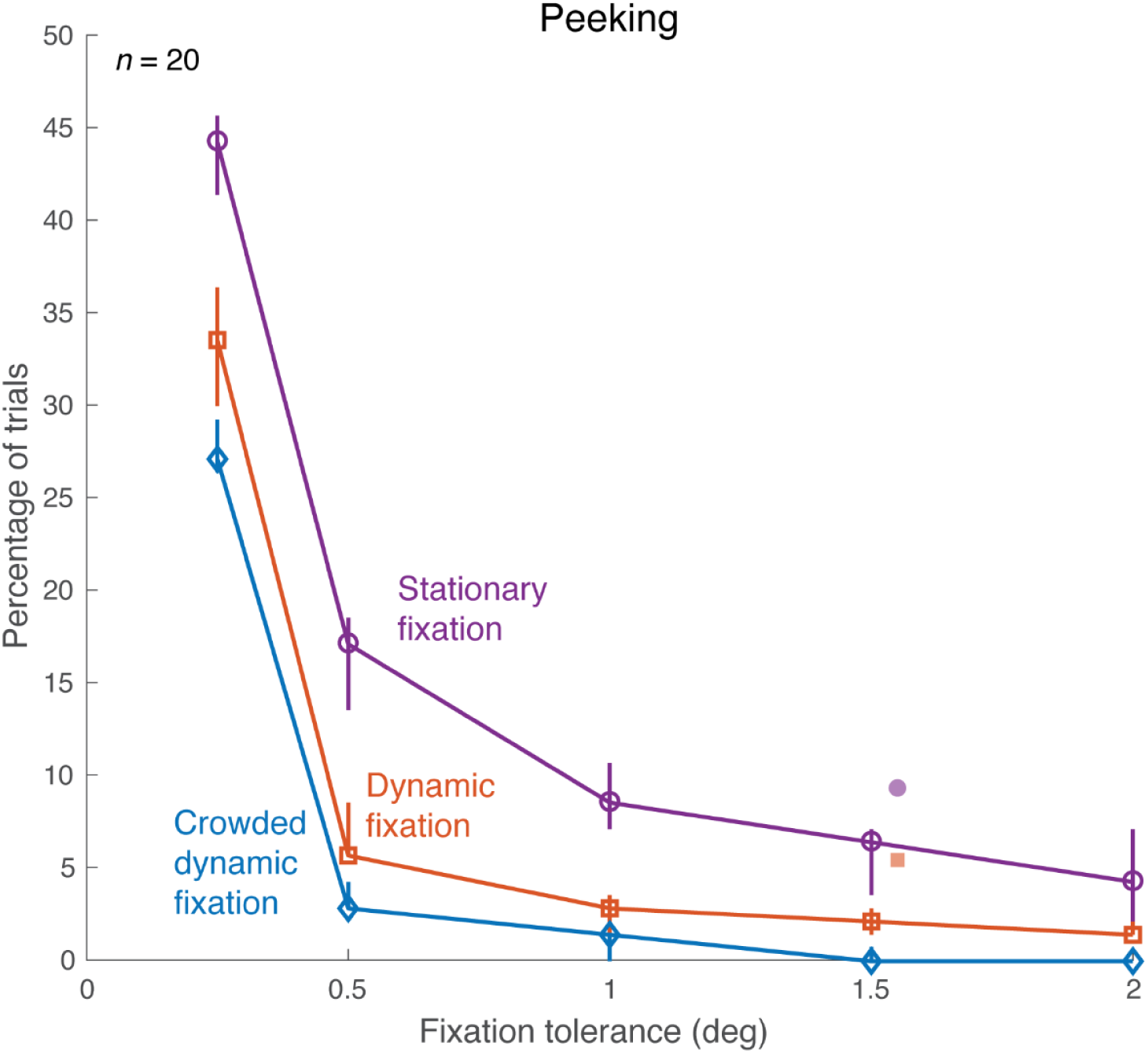
Frequency of fixation breaks. For each fixation condition, with several different fixation-error tolerances (x-axis), fixation breaks were counted and shown as a percentage of total trials (y-axis).

Each open symbol on the curve represents the median percentage of trials, calculated from 40 sessions, where participants broke fixation during triplet presentations. Error bars indicate 68% confidence intervals bootstrapped across sessions. Filled symbols: Kurzawski et al. (2023), horizontally offset by 0.05 deg for display purposes. *(plot_peek_number_sweep.m* and *plot_peeking_summaries.m)*.

### Crowded Dynamic Tracking eliminates abnormally small crowding thresholds that can be caused by peeking

The main goal of this study is improved fixation. Here we consider how the improved fixation affected the measured peripheral thresholds. Figure 6 summarizes crowding thresholds estimated with each fixation task. The key in this figure is the reduced range of the crowding thresholds in the crowded dynamic condition (Figure 6A). Kurzawski, Burchell, and colleagues (2023) measured Bouma factors of 50 participants with and without a gaze-contingent procedure, and found that the gaze-contingent procedure increased the minimal Bouma factor. Among the 200 Bouma factors the authors obtained at 10 deg eccentricity on the left and right horizontal meridians (50 participants, tested twice on each meridian), the minimal Bouma factor was 0.096. We take this Bouma factor as the minimum expected Bouma factor when there is no gaze error (i.e. no “peeking”). We find that with stationary and dynamic fixation, around 7% of the crowding thresholds are below the expected minimum, presumably due to peeking (Figure 6A, top and middle panels). Crowded dynamic fixation greatly reduces peeking (Figure 5) and abolishes these abnormally small thresholds (Figure 6A, bottom panel). The abnormally small thresholds in the stationary and dynamic fixation conditions have a small effect on the estimated average Bouma factors (Figure 6B). Examining only the 5 out of 36 experimental sessions that had no fixation breaks (peeking), the crowding thresholds do not differ across the three fixation conditions (Figure 6C). In all three conditions, we observe the expected left-right asymmetry, confirming prior findings of a right-field advantage (Kurzawski, Burchell, et al., 2023).

**Figure 6.**
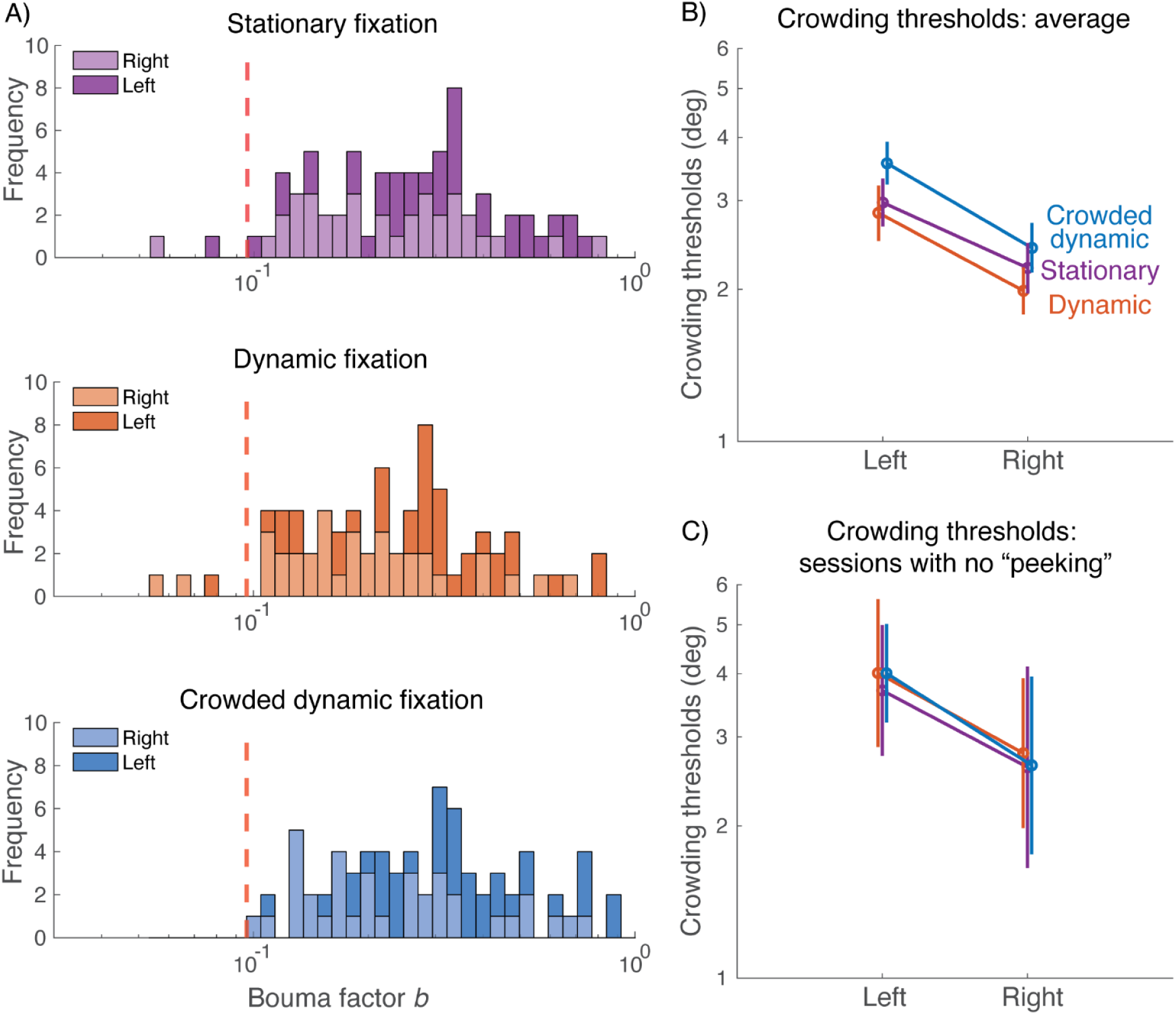
Crowding thresholds measured with the three fixation tasks. A) Stacked histograms of all Bouma factors acquired under each fixation condition (n = 72 for each condition, 18 observers x 2 locations x 2 repeats). Bouma factors to the left of the red dashed line (*b* = 0.096) are considered abnormally small *(plot_thresholds.m)*. B). Geometric mean of crowding thresholds for 19 participants *(plot_thresholds.m)*. C) Geometric mean of crowding thresholds for the 5 sessions (3 participants) with no fixation breaks (tolerance = 1.5 deg) in any condition. For panels B and C, the data points are jittered horizontally for visibility *(plot_thresholds_obedient_participants.m)*.

### With crowded dynamic fixation, manual tracking is difficult while averting gaze

Distractors were included in crowded dynamic fixation so that gaze aversion would result in crowding that would challenge manual tracking. We verified that gaze aversion worsens manual tracking by asking 4 participants to manually track the crosshair while fixating on a red dot that was 1, 2, 4, or 8 deg displaced from the crosshair (Figure 7A). When participants fixated farther away from the crosshair, cursor tracking error increased, and it increased more if the distractors were present (crowded dynamic fixation) than if the distractors were absent (dynamic fixation) (Figure 7B). The inset to panel B shows an example of the large decline in manual tracking accuracy when fixating a dot 8 deg away from the crosshair. In addition, compared to dynamic fixation, participants reported feeling more challenged with crowded dynamic tracking when required to look away (Figure 7C). However, when participants were foveating the crosshair, cursor tracking error was equally low for both dynamic and crowded dynamic tracking, as was perceived difficulty. This shows that adding the distractors does not add tracking difficulty when participants maintain central fixation. This is expected for crowding, which is generally negligible at fixation and grows linearly with eccentricity (Bouma, 1971; Rosen et al., 2014).

**Figure 7.**
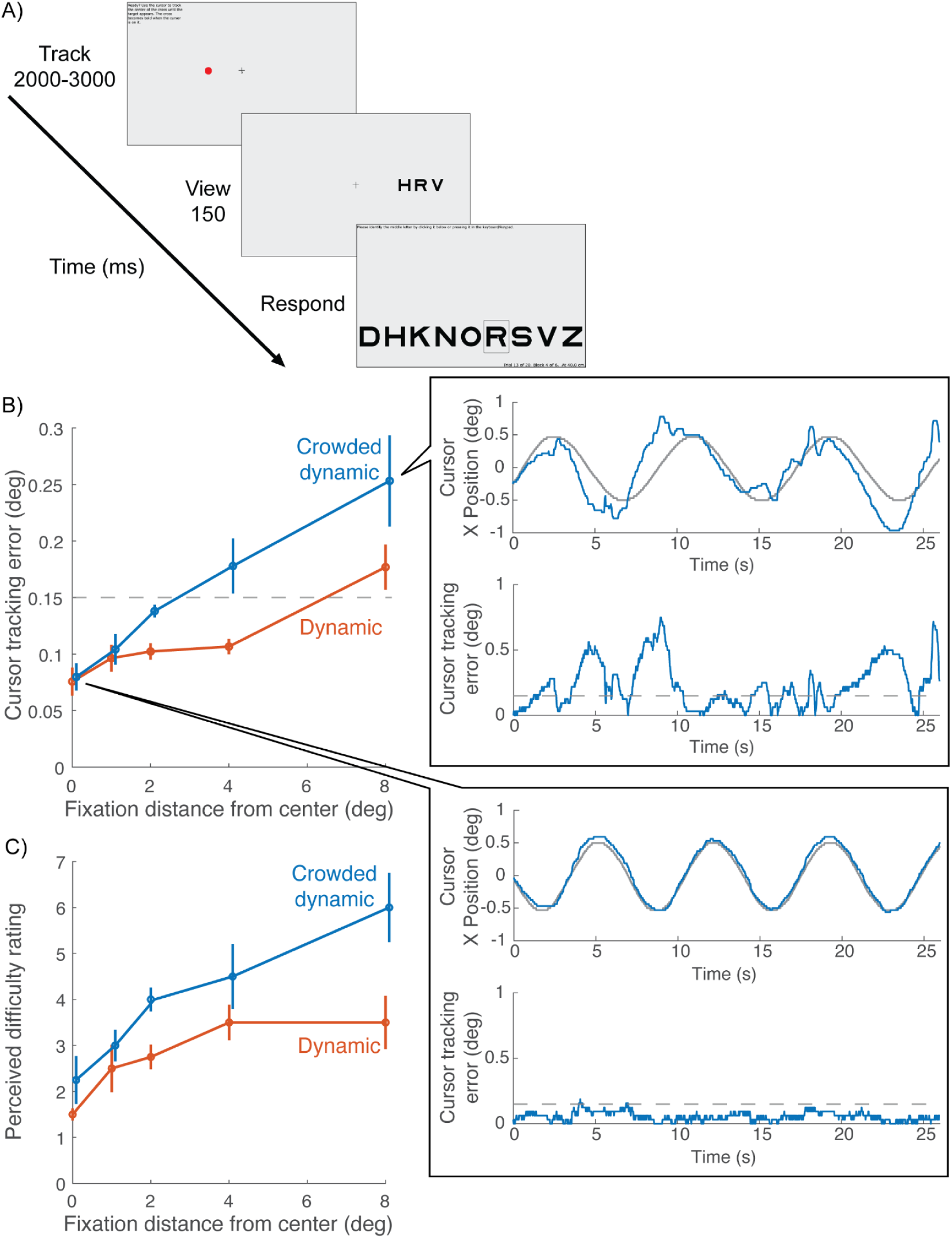
Cursor tracking accuracy while fixating elsewhere (N = 4). The purpose was to determine how well participants could track peripherally. A) The fixation task was either Dynamic or Crowded Dynamic (Figure 1). In both cases, participants tracked the moving crosshair with the cursor. While doing so, they either fixated the crosshair (0 distance) or a red dot that was 1, 2, 4, or 8 deg displaced from the crosshair. B) Cursor tracking error as a function of the distance between fixation and the crosshair *(flies_noFlies_dot.m)*. The blue curve is shifted horizontally to avoid occlusion. Blowout plots cursor tracking data from an example participant in the Crowded dynamic condition when the person fixated 8 deg from the screen center (top blowout) and fixate at the crosshair (bottom blowout). The top figure of each blowout plots X positions of cursor (blue curve) and crosshair positions (gray curve) over time. The bottom figure of each blowout plots cursor tracking error over time (blue curve) and the dashed gray line marks the 0.15-deg hotspot radius (*plot_tracking_demo.m*). C) Perceived difficulty rating for the same task as B *(flies_noFlies_rating.m)*.

### Crowded dynamic fixation slightly increases experiment duration

The two dynamic fixation tasks slightly lengthen the experiment because they add a tracking interval to each trial prior to stimulus onset. Figure 8 shows the impact of the fixation task on several durations. Tracking duration per trial increases (Figure 8A), but viewing (150 ms, not shown) and response duration (Figure 8B) are unchanged, resulting in slightly increased duration of the 70-trial block (Figure 8C): 5.5 min for Stationary, 6.2 min for Dynamic, and 6.5 min for Crowded dynamic. Thus, an experiment with Crowded dynamic fixation costs 5% longer than one with Dynamic fixation, and 18% longer than one with Stationary fixation.

**Figure 8.**
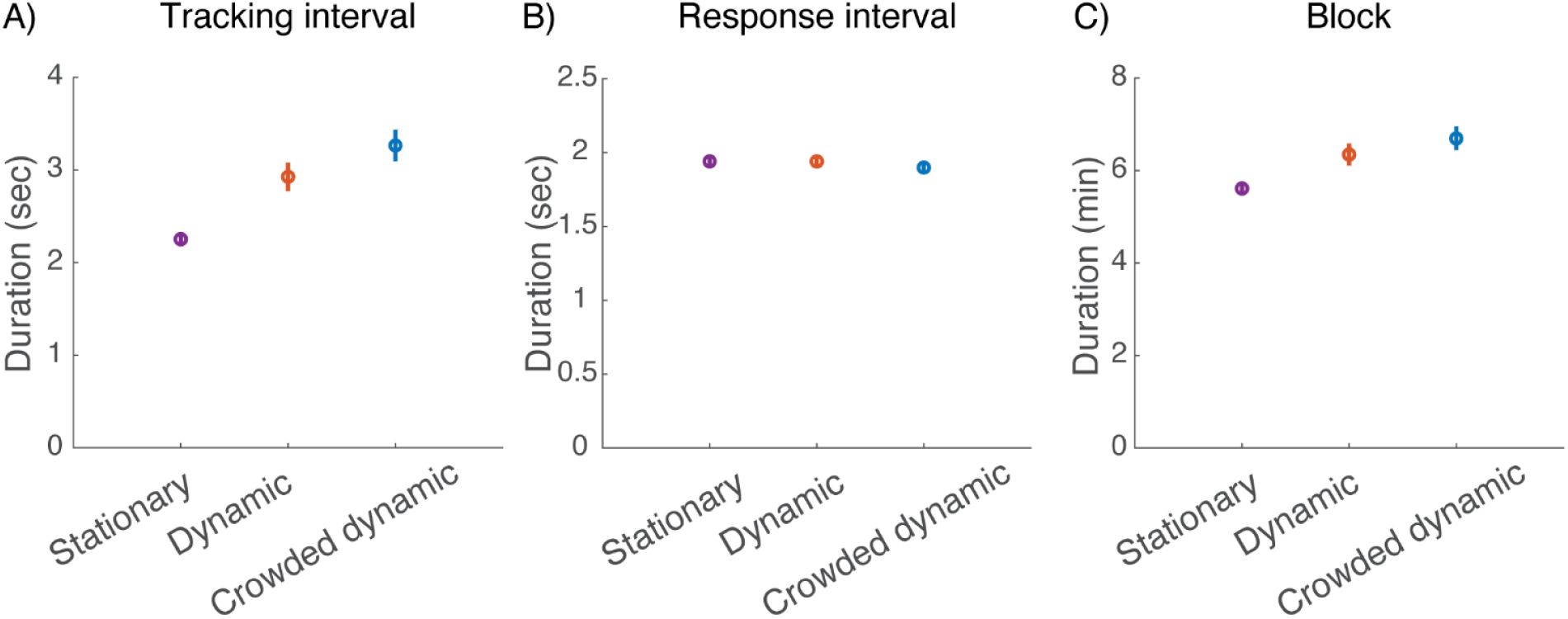
Effect of fixation task on experiment duration (20 participants, two sessions each). A) Average tracking interval duration. B) Average response interval duration. C) Average duration of 70-trial block. Error bars represent ±standard error of the mean. (*plot_time_spent.m*).

## Discussion

The current study compared the accuracy of stationary fixation, dynamic fixation, and crowded dynamic fixation while participants performed a peripheral task. Gaze accuracy increased from stationary to dynamic to crowded dynamic fixation. The reliable gaze accuracy of crowded dynamic fixation improved our estimates of threshold on the peripheral task, eliminating the abnormally small thresholds observed with the other two fixation tasks. In other words, crowding dynamic fixation greatly reduces the frequency of peeking, and abolishes the abnormally low thresholds that are caused by peeking.

### Why does cursor tracking improve fixation?

We instructed participants to manually track the moving crosshair, hoping that they would track it with their eyes. We need the cursor task because we can track the cursor, but not gaze, online. It is natural to look at an object when manually tracking it. Indeed, experiments measuring hand and eye movements show that they are highly correlated during target tracking (Koken & Erkelens, 1992; Xia & Barnes, 1999). Our in-lab EyeLink ground truth measurements showed that this works very well, confirming the results from Kurzawski, Pombo et al. (2023).

### The importance of fixation

Accurate knowledge of gaze position, whether by gaze tracking or accurate fixation, is essential for study of many visual tasks. An obvious example is peripheral thresholds, which typically depend strongly on eccentricity. For example, from 0° to 10° eccentricity, grating acuity threshold (deg per cycle) increases about 4-fold (Frisén & Glansholm, 1975), letter acuity threshold (minimal angle of resolution) increases about 7-fold (Ludvigh, 1941), and crowding distance increases about 40 fold (Kurzawski, Burchell, et al., 2023). Inconsistent fixation, “peeking,” produces misleading results: When participants shift their gaze towards a peripheral target, estimated thresholds are profoundly lower (Kurzawski, Burchell, et al., 2023).

Location matters for other reasons too. For example, fixation control is essential in adaptation experiments, where prolonged exposure to a stimulus at a specific visual field location affects perception most strongly at that location (Gibson, 1937). Many studies of object recognition (Potter, 1976) and scene perception (Fei-Fei et al., 2007) ask how much information can be processed from a single glance; in such studies it is important to know where the participant fixates. In attention experiments, a peripheral cue improves processing for subsequent stimuli in nearby (but not distant) locations (Carrasco, 2011; Posner, 1980), hence it is important that participants do not move their eyes during the interval between presentations of cue and target. Similarly, in change-blindness experiments, it is important to know whether participants maintain central fixation or move their eyes to the location of the image where there is a change (Rensink et al., 1997; Simons & Rensink, 2005).

### The webcam alternative

Another way to ensure fixation during online vision experiments is webcam-based eye tracking. Saxena et al. (2022) and Papoutsaki et al. (2016) reported that WebGazer, an open-source webcam-based eye-tracking tool, measures gaze to within 4 deg. This is useful for experiments that need dynamic gaze tracking, but is typically not precise enough to ensure fixation for threshold testing. Kaduk et al. (2024) showed that one can improve the gaze precision to 1.5 deg by adding an external high-resolution webcam. For reference, the gold standard for lab-based vision studies is EyeLink 1000 with a precision of 0.5 deg. However, adding an external webcam limits the study to participants willing to obtain and install additional equipment.

Because our tracking task does not require equipment beyond a typical computer, it can be used at scale for online testing. The experiments presented in this paper were conducted in the lab in order to use the EyeLink for ground truth, but with our online web-based software. In addition, previous work by Kurzawski, Pombo et al. (2023) demonstrated that in-lab and online testing with EasyEyes yield consistent results.

### Does the crowded dynamic fixation task draw more attention than a static fixation mark?

To some extent, typical peripheral testing is always a dual task, because the participant needs to direct their gaze to the fixation mark while anticipating a peripheral target (Pashler, 1994). Does moving and cluttering the fixation mark make the fixation task draw more attention away from the target? We do not have an assessment of the effect of motion on attention, but we did ask four participants to rate the perceived difficulty of tracking the moving crosshair with and without distractors. Admittedly, effort and attention are not quite the same thing. The rated effort was similar for dynamic and crowded dynamic fixation when participants were fixating the crosshair.

We compared crowding thresholds measured with the three fixation tasks. We saw in Figure 6 that crowding thresholds differ little among the three fixation tasks. In order not to confound peeking with attention, we re-analyzed those data including only participants who did not peek in any condition (keeping 3 out of 20). Figure 6C, again, shows little difference between thresholds measured with the three fixation tasks. Thus, the crowding thresholds do not differ in a way that could be attributed to differing demands of the three fixation tasks. It would be interesting to assay this again with a task that is more sensitive to attention.

## Conclusions

Kurzawski, Pombo, and others (2023) introduced a manual tracking task for gaze control in psychophysical testing. We improved it by adding independently moving distractors, ensuring gaze remains on the crosshair. Fixation accuracy was verified during peripheral threshold measurement. This task requires no extra equipment and suits online testing. Crowded dynamic fixation provides gaze control for online visual testing, facilitating larger and more diverse samples with minimal effort.

## Acknowledgements

We thank Jan Kurzawski (U Maastricht) and Maria Pombo (NYU) for helping us improve their gaze control method. Funding support from NIH grants R01-EY027401 (JW), R01-EY027964 (DGP) and R01-EY033628 (JW).

## Data and code availability

Raw data and code used to analyze the data reside in https://github.com/fh986/crowded-dynamic-fixation.

## Supplementary figures

**Supplement 1.**
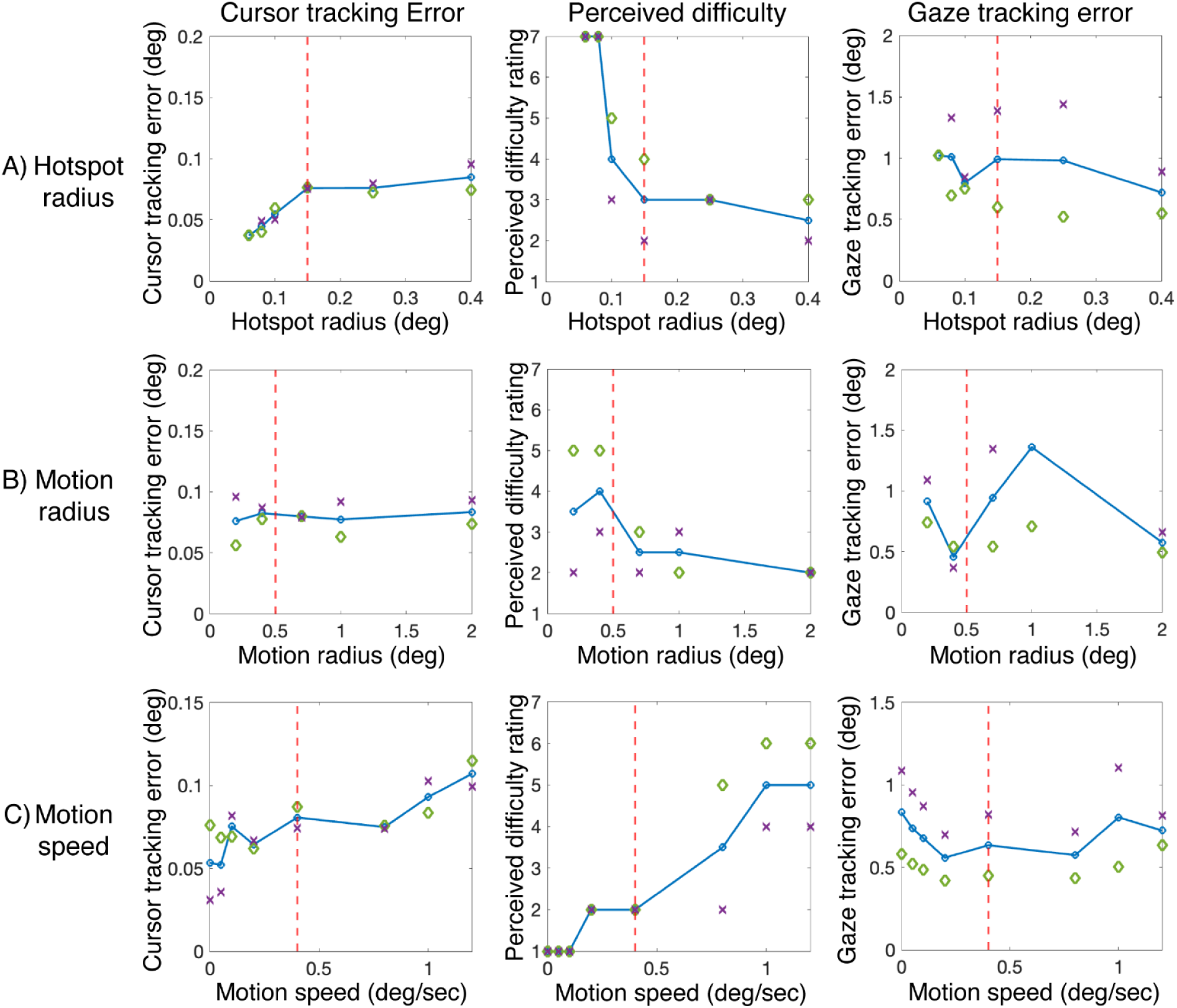
Exploratory mapping of crosshair parameters, for adults. Vertical red dashed lines indicate the parameter values used in this paper. A) Hotspot radius: A radius smaller than 0.15 deg is too challenging, but a hotspot radius of 0.15 deg is moderately challenging and keeps the cursor and gaze tracking errors low; B) Motion radius: A motion radius below 0.4 deg is too challenging, and a radius larger than 0.4 deg produces too much gaze tracking error; C) Motion speed: Speeds below 0.2 deg/sec are too easy, and speeds much above 0.4 deg/sec are too challenging. *(compare_ee_el_wTimeouts.m)*

